# Melissa: Bayesian clustering and imputation of single cell methylomes

**DOI:** 10.1101/312025

**Authors:** Chantriolnt-Andreas Kapourani, Guido Sanguinetti

## Abstract

Measurements of DNA methylation at the single cell level are promising to revolutionise our understanding of epigenetic control of gene expression. Yet, intrinsic limitations of the technology result in very sparse coverage of CpG sites (around 5% to 20% coverage), effectively limiting the analysis repertoire to a semi-quantitative level. Here we introduce Melissa (MEthyLation Inference for Single cell Analysis), a Bayesian hierarchical method to quantify spatially-varying methylation profiles across genomic regions from single-cell bisulfite sequencing data (scBS-seq). Melissa clusters individual cells based on local methylation patterns, enabling the discovery of epigenetic differences and similarities among individual cells. The clustering also acts as an effective regularisation method for imputation of methylation on unassayed CpG sites, enabling transfer of information between individual cells. We show both on simulated and real data sets that Melissa provides accurate and biologically meaningful clusterings, and state-of-the-art imputation performance. An R implementation of Melissa is publicly available at https://github.com/andreaskapou/Melissa.

## 1 Introduction

DNA methylation is probably the best studied epigenomic mark, due to its well established heritability and widespread association with diseases and a broad range of biological processes, including X-chromosome inactivation, cell differentiation and cancer progression (Baylin and Jones, 2011; Bird, 2002; Jones, 2012). Yet its role in gene regulation, and the molecular mechanisms underpinning its association with diseases, are still imperfectly understood.

Bisulfite treatment of DNA followed by sequencing (BS-seq) has provided a powerful tool for measuring the methylation level of cytosines on a genome-wide scale with single nucleotide resolution (Krueger *et al*., 2012). BS-seq protocols have been vastly improved over the last decade, with BS-seq rapidly becoming a widespread tool in biomedical investigation. Nevertheless, until very recently BS-seq could only be used to measure methylation in bulk populations of cells (Shapiro *et al*., 2013), preventing effective investigations of the role of DNA methylation in shaping transcriptional variability and early development (Kelsey *et al*., 2017; Schwartzman and Tanay, 2015).

This shortcoming has been addressed within the last five years through the development of protocols to measure DNA methylation at single-cell resolution using either scBS-seq (Smallwood *et al*., 2014) or scRRBS (Guo *et al*., 2013) making it possible to uncover the heterogeneity and dynamics of DNA methylation (Farlik *et al*., 2015). Even more recently, methods have been developed that can sequence both the methylome and the transcriptome or other features in parallel, potentially enabling a quantification of the role of DNA methylation in explaining transcriptional heterogeneity (Angermueller *et al*., 2016; Clark *et al*., 2018; Hou *et al*., 2016). However, due to the small amounts of genomic DNA per cell, these protocols usually result in very sparse genome-wide CpG coverage, ranging from 5% in high throughput studies (Luo *et al*., 2017; Mulqueen *et al*., 2018) to 20% in low throughput ones (Angermueller *et al*., 2016; Smallwood *et al*., 2014). The sparsity of the data represents a major hurdle to effectively use single-cell methylation data to inform our understanding of epigenetic control of transcriptomic variability, or to distinguish individual cells based on their epigenomic state.

In this paper, we address these problems by using a two-pronged strategy. First, we note that several recent studies have highlighted the importance of local methylation profiles, as opposed to individual CpG methylation, in determining the epigenetic state of a locus (Kapourani and Sanguinetti, 2016; Mayo *et al*., 2015; Vanderkraats *et al*., 2013). This implies that local spatial correlations may be effectively leveraged to ameliorate the issue of data sparsity. Secondly, single-cell BS-seq protocols, as all single-cell high-throughput protocols, simultaneously assay a large number of cells, ranging from several tens (Smallwood *et al*., 2014) to a few thousands in the most recent studies (Luo *et al*., 2017). Such abundance of data could be exploited to our advantage to transfer information across similar cells.

We implement both of these strategies within Melissa (MEthyLation Inference for Single cell Analysis), a Bayesian hierarchical model that jointly learns the methylation profiles of genomic regions of interest and clusters cells based on their genome-wide methylation patterns. In this way, Melissa can effectively use both the information of neighbouring CpGs and of other cells with similar methylation patterns in order to predict CpG methylation states. As an additional benefit, Melissa also provides a Bayesian clustering approach capable of identifying subsets of cells based solely on epigenetic state, to our knowledge the first clustering method tailored specifically to this rapidly expanding technology. We benchmark Melissa on both simulated and real single-cell BS-seq data, demonstrating that Melissa provides both state of the art imputation performance, and accurate clustering of cells. Furthermore, thanks to a fast variational Bayes estimation strategy, Melissa has good scalability and can provide an effective modelling tool for the increasingly large single-cell methylation studies which will become prevalent in coming years.

## 2 Results

Melissa addresses the data sparsity issue by leveraging local correlations and similarity between individual cells (see Fig. 1). The starting point is the definition of a set of genomic loci or regions (e.g. genes or enhancers). Within each locus, Melissa postulates a latent profile of methylation, a function mapping each CpG within the locus to a number in [0,1] which defines the probability of that CpG being methylated. To ensure spatial smoothness of the profile, Melissa uses a generalised linear model of basis function regression along the lines of Kapourani and Sanguinetti (2016) (with modified likelihood to account for single cell data). Local correlations are however often insufficient for loci with extremely sparse coverage, and these are quite common in scBS-seq data. Therefore, we share information across different cells by coupling the local GLM regressions through a shared prior distribution. In order to respect the (generally unknown) population structure that may be present within the cells assayed, we choose a (finite) Dirichlet mixture model prior.

**Figure 1:**
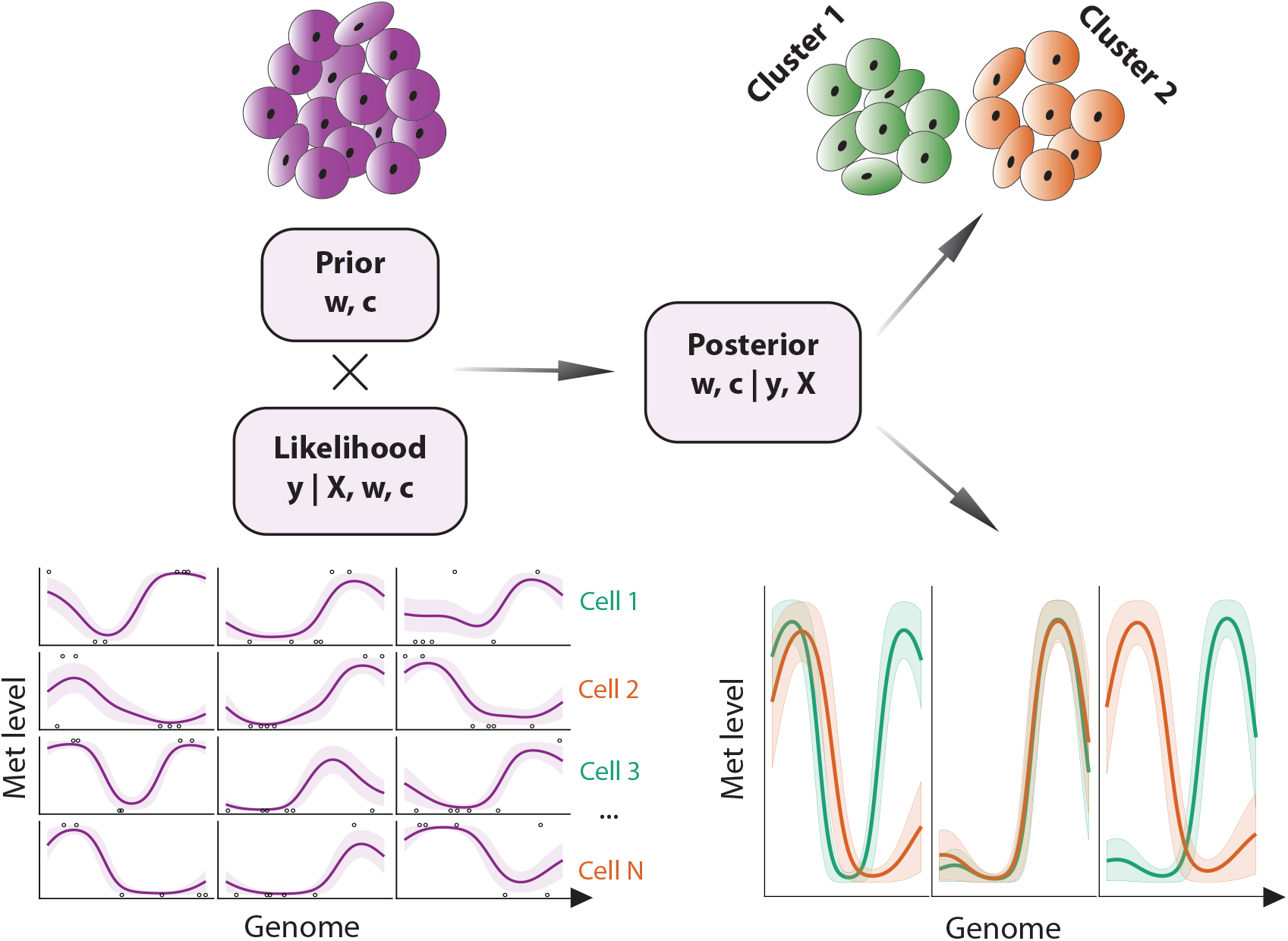
Melissa model overview. Melissa combines a likelihood computed from single cell methylation profiles (bottom left) and a Bayesian clustering prior (top left). The posterior distribution provides a methylome-based clustering (top right) and imputation (bottom right) of single cells.

The output of Melissa is therefore twofold: at each locus in each cell, we get a predicted profile of methylation, which can be used to impute missing data. For each cell, we also get a discrete cluster membership probability, providing a methylome-based clustering of cells.

### 2.1 Benchmarking Melissa on simulated data

We benchmark the ability of our model to cluster and impute CpG methylation states at the single cell level both on simulated and mouse embryonic stem cell (ESC) data sets. To assess test prediction performance we consider different metrics, including F-measure, the area under the receiver operating characteristics curve (AUC) and precision recall curves (Powers, 2011).

To benchmark the performance of *Melissa* in predicting CpG methylation states, we compare it against five different imputation strategies. As a baseline approach, we compute the average methylation rate separately for each cell and region (*Indep Rate*). We also use the BPRMeth model (Kapourani and Sanguinetti, 2016), where we account for the binary nature of the observations, which we train independently across cells and regions (*Indep Profile*). To share information across cells, but not across neighbouring CpGs, we constrain Melissa to infer constant functions, i.e. learn average methylation rate (*Melissa rate*). Additionally, as a fully independent baseline, we use a Random Forest classifier trained on individual cells and regions using the methylation state of covered CpGs as input features (*RF*); this is essentially the method of Zhang et al. (2015), but without using additional annotation data or DNA sequence patterns. We delay comparisons with the deep learning method DeepCpG (Angermueller *et al*., 2017) to the next section, as DeepCpG is not applicable in the settings of our simulation (see later Section 2.3).

In order to generate realistic simulated single-cell DNA methylation data, we extracted methylation profiles from real (bulk) BS-seq data using the BPRMeth package (Kapourani and Sanguinetti, 2018), subsequently downsampling to simulate the low coverage of scBS-seq. In total we simulated *N* = 200 cells from *K* = 4 sub-populations, where each cell consisted of *M* = 100 genomic regions. Additionally, to account for different levels of similarity between cell sub-populations, we simulated 11 different datasets by varying the proportion of similar genomic regions between clusters (see “Methods” section).

Applying the competing methods to synthetic data we observe that *Melissa* yields a substantial improvement in prediction accuracy compared to all other models (Fig. 2, Additional file 1: Fig. S1). Notably, *Melissa* is robust across different settings of the data, such as variable CpG coverage in each region (Fig. 2a), or different levels of dissimilarity across clusters (Fig. 2b). Due to its ability to transfer information across cells and neighbouring CpGs, our model robustly maintains its prediction accuracy even at a very sparse coverage level of 10%. The *Indep Profile* and *RF* models perform poorly at low CpG coverage settings, becoming comparable to *Melissa* when using the majority of the CpGs for training set. Importantly, *Melissa* still performs better at 90% CpG coverage, demonstrating that the clustering acts as an effective regularisation for imputing unassayed CpG sites. As expected, *Indep Rate* and *Melissa Rate* methods had a poor performance, since they are not expressive enough to capture spatial correlations between CpGs. The imputation performance of all methods is relatively insensitive to the degree of cluster dissimilarity (Fig. 2b).

**Figure 2:**
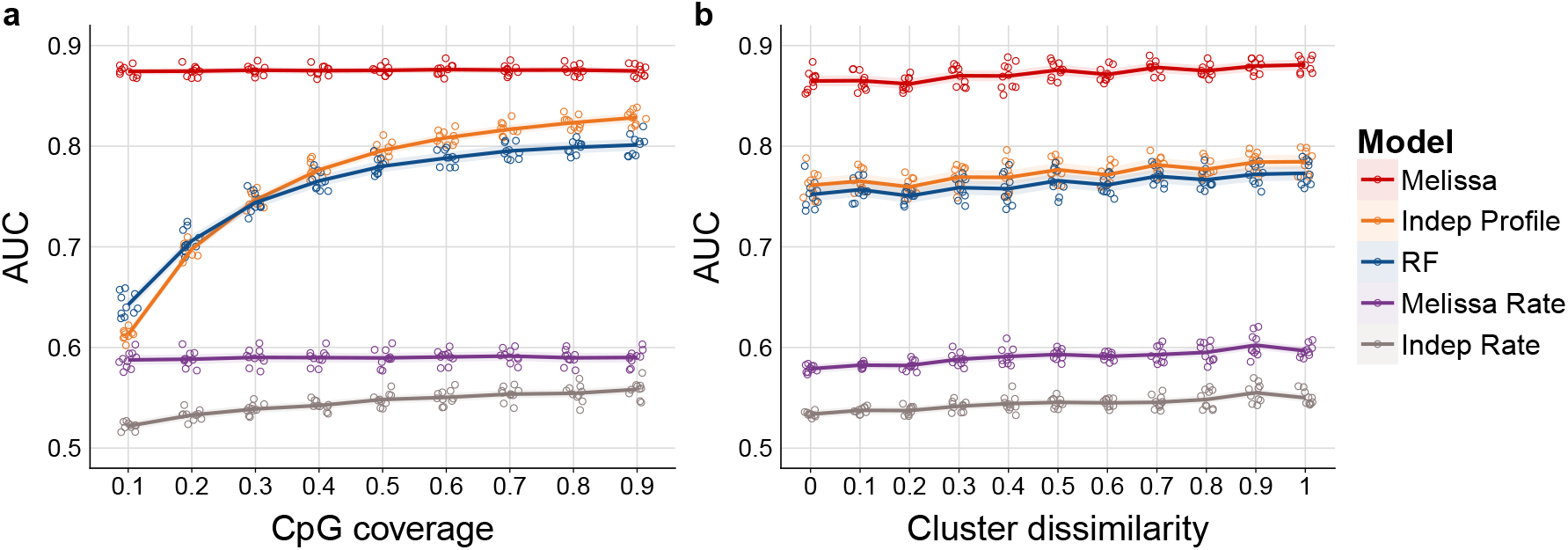
Melissa robustly imputes CpG methylation states. (**a**) Imputation performance in terms of AUC as we vary the proportion of covered CpGs used for training. Higher values correspond to better imputation performance. For each CpG coverage setting a total of 10 random splits of the data to training and test sets was performed. Each coloured circle corresponds to a different simulation. The plot shows also the LOESS curve for each method as we increase CpG coverage. (**b**) Imputation performance measured by AUC for varying proportions of similar genomic regions between clusters. Values closer to zero correspond to highly similar cell sub-populations, whereas values closer to one correspond to well separated cell sub-populations. In (**a**) cluster dissimilarity was set to 0.5 and in (**b**) CpG coverage was set to 0.4.

Next we consider the clustering performance of *Melissa*. Since most of the rival methods do not have a notion of clustering, we compare *Melissa* to clustering using methylation rates (*Melissa Rate*). As a performance metric, we use the Adjusted Rand Index (ARI) (Hubert and Arabie, 1985) between the true cluster assignment and the predicted cluster membership returned from the model. Fig. 3a shows ARI values comparing the two models for varying CpG coverage (with cluster dissimilarity level at 0.5). *Melissa* performs perfectly in all settings, demonstrating its power and sensitivity in identifying robustly the cell sub-population structure. When varying the level of cluster dissimilarity (see Fig. 3b), the model is still able to retain its high clustering performance. As expected, for settings with low variability between clusters (i.e. cell sub-populations are difficult to distinguish), the performance drops; however, *Melissa* is consistently superior to the *Melissa Rate* model, and rapidly reaches near-perfect clustering accuracy.

**Figure 3:**
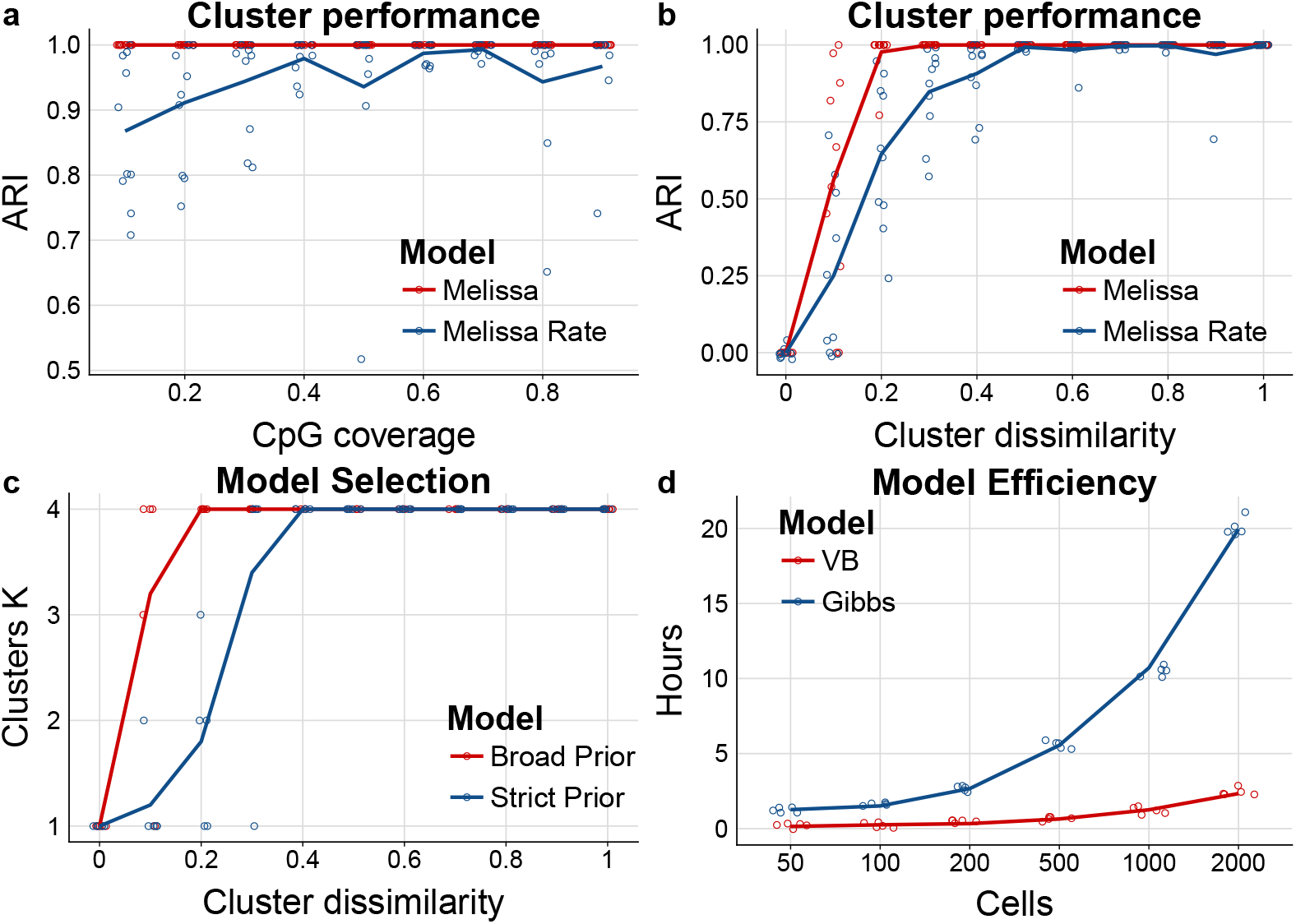
Melissa efficiently and accurately clusters cell sub-populations. (**a**) Clustering performance measured by ARI as we vary CpG coverage. Higher values correspond to better agreement between predicted and true cluster assignments. For each CpG coverage setting a total of 10 random splits of the data to training and test sets was performed. Each coloured circle corresponds to a different simulation. The plot shows also the LOESS curve for each method as we increase CpG coverage. (**b**) Clustering performance (ARI) for varying proportions of similar genomic regions between clusters. (**c**) Predicted number of clusters using two different prior settings: a broad and a strict prior as we vary cluster dissimilarity. Initial number of clusters was set to *K* = 10. Melissa identifies the correct number of clusters in most parameter settings (*K* = 4); notably when there is no dissimilarity across clusters (i.e. we have one global cell sub-population), Melissa prunes away all components and keeps only one cluster (*K* = 1). (**d**) Running times for varying number of cells for the variational Bayes (VB) and Gibbs sampling implementations for the Melissa model, where each cell consists of *M* = 200 genomic regions.

Subsequently, we test *Melissa*’s ability to perform model selection, that is to identify the appropriate number of cell sub-populations. To do so, we run the model on simulated data, setting the initial number of clusters to *K* = 10 and letting the variational optimisation prune away inactive clusters (Corduneanu and Bishop, 2001). We used both broad (red line) and shrinkage (blue line) priors. Fig. 3c shows that the variational optimisation automatically recovered the correct number of mixture components for almost all parameter settings. As expected, in settings with high between cluster similarity, the model with shrinkage prior returned fewer clusters, since the data complexity term in Eq. (13) (see “Methods” section) was penalizing more the variational approximation compared to the gain in likelihood from explaining the data.

Finally, we assess the scalability of *Melissa* with respect to the number of single cells. Fig. 3d compares the variational Bayes (red line) with the Gibbs sampling (blue line) algorithm; which demonstrates the good scalability of VB where we can analyse thousands of single cells in acceptable running times. The maximum number of iterations for the VB algorithm was set to 400 and the Gibbs algorithm was run for 3000 iterations. Both algorithms are implemented in the R programming language and were run on a machine utilising at most 16 CPU cores.

### 2.2 Melissa accurately predicts methylation states on real data

To assess Melissa’s performance on real scBS-seq data we considered two mouse embryonic stem cell (ESC) data sets from Angermueller et al. (2016) and Smallwood et al. (2014). The mouse ESCs were cultured either in 2i medium (*2i ESCs*) or serum conditions (*serum ESCs*), hence we expect methylation heterogeneity between cell sub-populations. In addition, in *serum ESCs* there is evidence of additional CpG methylation heterogeneity (Ficz *et al*., 2013) making these data suitable for the model selection task to infer cell sub-population numbers. The analysis on both data sets was performed on six different genomic contexts: protein coding promoters with varying genomic windows: ±1.5kb, ±2.5kb and ±5kb around transcription start sites (TSS), active enhancers, super enhancers and Nanog regulatory regions (see “Methods” section for details on data preprocessing). In this section, we additionally compare *Melissa* to the deep learning model DeepCpG (Angermueller *et al*., 2017) that uses the information of neighbouring CpGs to predict the methylation state of each target CpG site. It should be noted that *DeepCpG* is designed to predict individual missing CpGs, rather than missing regions, and requires always information about neighbouring CpGs. This means that, during prediction, *DeepCpG* always has access to more data than competing methods, potentially providing it with an unfair advantage; to partly address this problem, we also present results when DeepCpG had access to subsampled data (labelled *DeepCpG Sub* in our figures). In general, *DeepCpG* should be thought as complementary to Melissa, and comparisons should be evaluated cautiously (see below Section 2.3).

We first applied *Melissa* on the Angermueller et al. (2016) data set which consists of 75 single cells (14 *2i ESCs* and 61 *serum ESCs*). Fig. 4a shows a direct comparison of the imputation performance of all the methods across a variety of genomic contexts. *Melissa* is better or comparable to rival methods in terms of AUC (see Fig. 4a), and substantially more accurate in terms of F-measure (Additional file 1: Fig. S2), demonstrating its ability to capture local CpG methylation patterns. *DeepCpG* also performs strongly on most genomic regions, indicating that a flexible deep learning method is effective in capturing patterns of methylation. Similar results were obtained by considering different metrics (Additional file 1: Fig. S2 - S4). Boxplots show performance distributions across 10 independent training / test splits of the data, except for *DeepCpG*, where the high computational costs prevented such investigation. Interestingly, methods based on methylation rates performed poorly at promoters, underlining the importance of methylation profiles in distinguishing epigenetic state near transcription start sites and identifying meaningful cell sub-populations. For all models, the imputation performance (in terms of AUC) at active enhancers was lower, indicating high methylation variability across cells and nearby CpG sites as shown in Smallwood et al. (2014).

**Figure 4:**
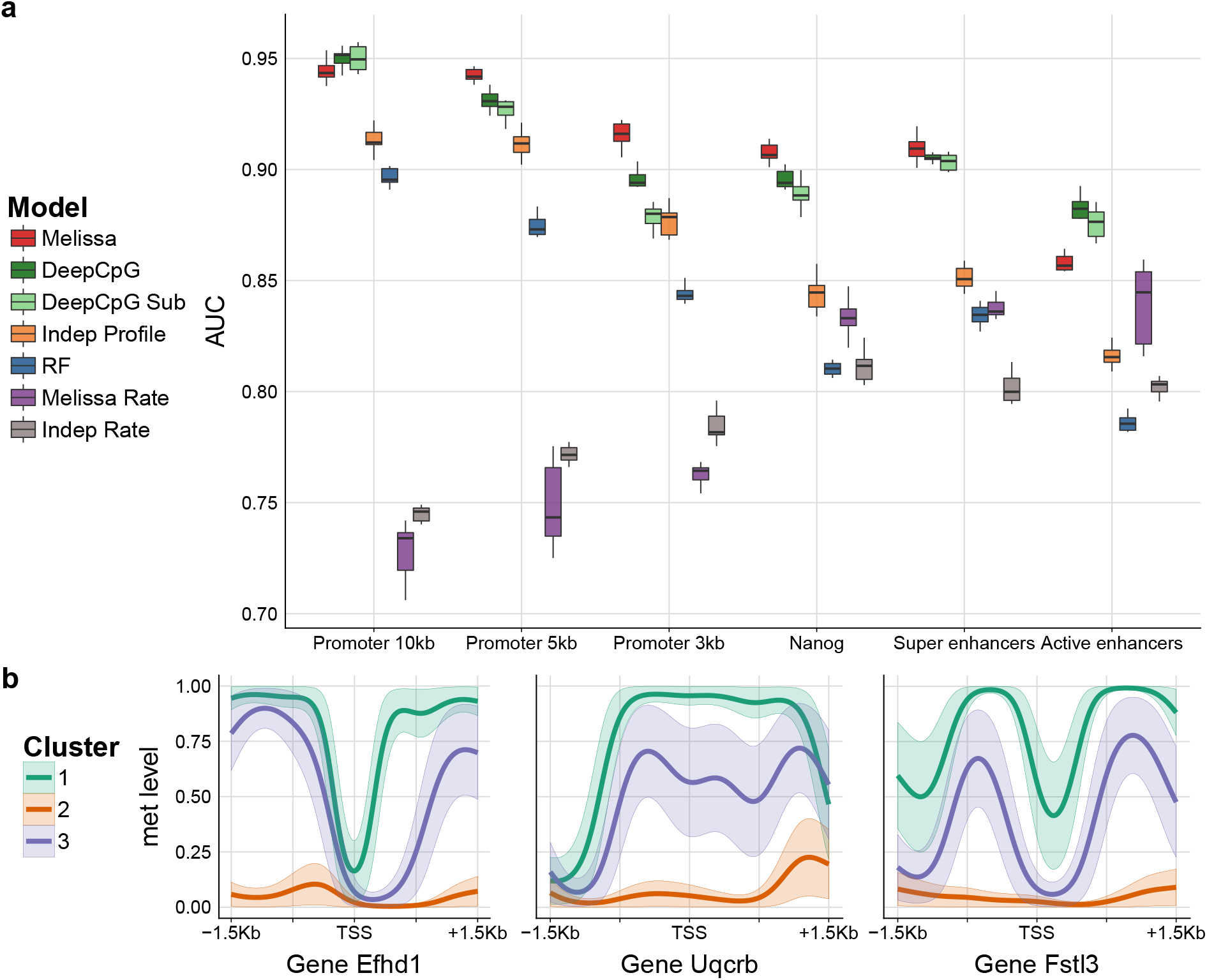
Imputation performance and clustering of mouse ESCs (Angermueller *et al*., 2016) based on genome wide methylation profiles. (**a**) Prediction performance on test set for imputing CpG methylation states in terms of AUC. Higher values correspond to better imputation performance. Each coloured boxplot indicates the performance using 10 random splits of the data in training and test sets; due to high computational costs, DeepCpG was trained only once and the boxplots denote the variability across ten random subsamplings of the test set. (**b**) Three example promoter regions with the predicted methylation profiles for the the Efhd1, Uqcrb and Fstl3 genes. Each coloured profile corresponds to the average methylation pattern of the cells assigned to each sub-population, in our case Melissa identified *K* = 3 clusters.

In terms of clustering performance, *Melissa* confirms that the data supports the existence of a sub-population of serum cells as suggested in Ficz et al. (2013), by returning three clusters in most contexts. However, for larger regions around promoters (±5kb and ±10kb) the model identified only two clusters, suggesting that the epigenetic differences between serum cell populations are subtle and that regulatory regions have higher epigenetic heterogeneity compared to promoters. Fig. 4b shows example promoter regions for three genes with their corresponding inferred methylation profiles for each cell sub-population. Each colour corresponds to a different cell sub-population, where orange-like profiles clearly correspond to *2i ESCs* which are globally hypo-methylated. Interestingly, 2i cells can be easily separated from serum cells based on methylation rate alone, due to the global hypo-methylation of 2i cells, however the sub-population structure within serum cells appears to be determined by changes in profiles.

As a second real data set, we analysed the smaller Smallwood et al. (2014) data set which consists of only 32 cells (12 *2i ESCs* and 20 *serum ESCs*). The imputation performance in terms of AUC across genomic contexts is shown in Fig. 5. Melissa retains its high prediction accuracy and is comparable with DeepCpG across most contexts (see Additional file 1: Fig. S5 - S7 for performance on different metrics), even though the full DeepCpG model has slightly better performance on this data set. This suggests that the small number of cells in this data set did not allow an effective sharing of information. In terms of clustering performance, when considering promoter regions *Melissa* confidently detected the expected clusters, easily separating 2i and serum cells. On the remaining genomic contexts the model identifies three clusters, mostly due to higher heterogeneity of these regions compared to promoters and once again underlying the emergence of epigenomically distinct populations within serum cells (see Additional file 1: Fig. S8, S9 for example methylation profiles across genomic contexts).

**Figure 5:**
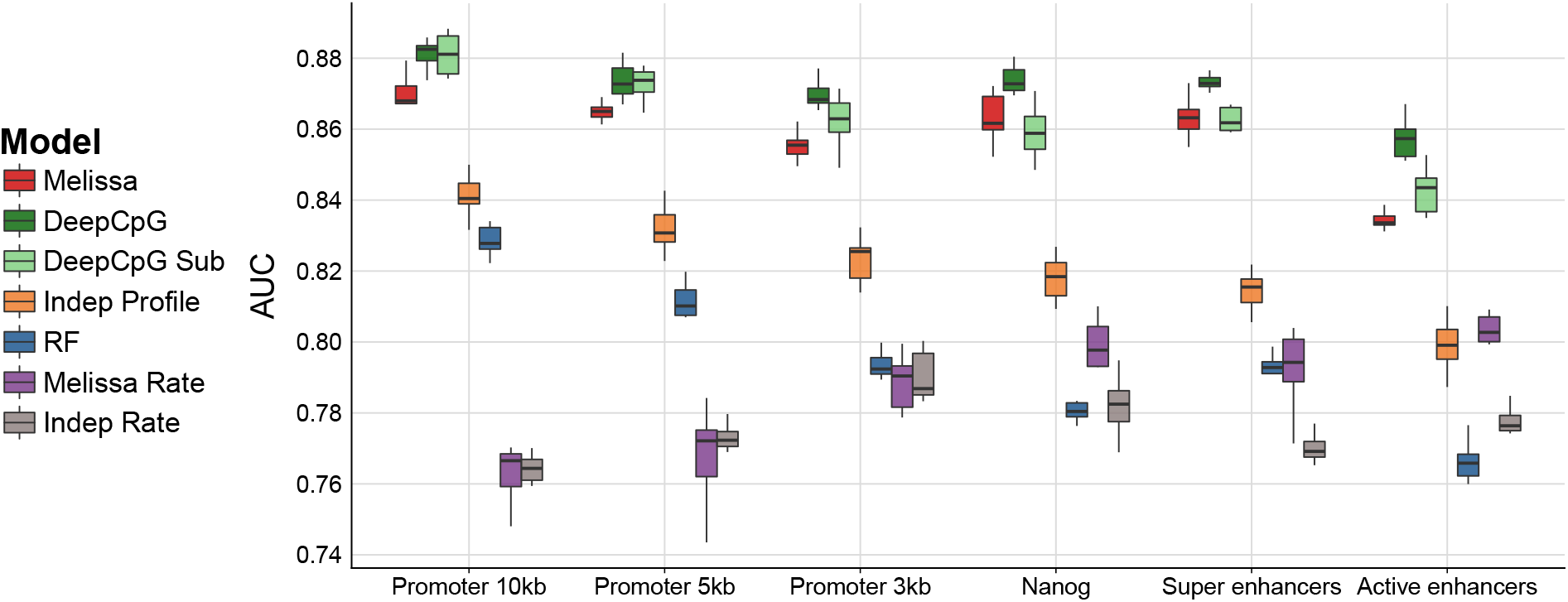
Imputation performance of mouse ESCs (Smallwood *et al*., 2014) based on genome wide methylation profiles. Shown is the prediction performance, in terms of AUC, for imputing CpG methylation states. Each coloured boxplot indicates the performance using 10 random splits of the data in training and test sets; due to high computational costs, DeepCpG was trained only once and the boxplots denote the variability across ten random subsamplings of the test set.

### 2.3 A note on the comparison with DeepCpG

Melissa and DeepCpG models reported substantially better imputation performance compared to the rival methods and show comparable performance when analysed on real datasets, demonstrating their flexibility in capturing complex patterns of methylation. However, the two methods have significantly different computational performances. In our experiments, Melissa’s runtime was less than seven hours for most genomic contexts running on a small server machine utilising at most ten CPU cores. By contrast, DeepCpG required around four days to analyse each dataset on a GPU cluster equipped with high end NVIDIA Tesla K40ms GPUs, and had very high memory requirements. These computational overheads effectively make DeepCpG out of reach for smaller research groups.

In addition, one should be cautious when directly comparing prediction performances due to the different design of the DeepCpG model. DeepCpG is trained on a specific set of chromosomes and considers each CpG site independently; hence it does not have a notion of genomic region to be trained on, and will in any case utilize information from neighbouring CpGs within or outside the region, information that Melissa and the rival methods do not have access to.

## 3 Discussion

Single cell DNA methylation measurements are rapidly becoming a major tool to understand epigenetic gene regulation in individual cells. Newer platforms are rapidly expanding the scope of the technology in terms of assaying large numbers of cells (Luo *et al*., 2017), however all technologies are plagued by intrinsically low coverage in terms of numbers of CpGs assayed.

In this paper, we have proposed Melissa as a way of addressing the low coverage issue by sharing information between CpGs with a local smoothing and between cells with a Bayesian clustering prior. On both synthetic and real data, Melissa achieved state of the art imputation performance over a panel of competing methods, including DeepCpG (Angermueller *et al*., 2017) and random forests. While achieving comparable or superior performance to black-box methods such as neural networks and random forests, Melissa is more transparent and needs minimal tuning: all the results shown, on both synthetic and real data, were obtained with the same settings of the algorithm. Additionally, as all Bayesian methods, Melissa outputs are probability distributions that fully quantify the uncertainty on the model’s prediction, and which are more easily usable for further experimental design compared to the point-estimates provided by black-box approaches. Melissa does not require additional annotation data as in Zhang *et al*. (2015) or Ernst and Kellis (2015), and does not exploit sequence information like DeepCpG, but an extension leveraging side data would be easily accomplished within the Bayesian framework and would represent an interesting extension for future research. By using a Bayesian clustering prior, Melissa has the added benefit of simultaneously uncovering the population structure within the assay, as we demonstrated in the real data examples; Melissa can therefore be a useful tool in uncovering epigenetic diversity among cells.

While Melissa accounts for heterogeneity in the cell population structure, it does not allow for heterogeneity at the single gene level: each cluster has a single methylation profile within each region, and all variability at the single locus level is attributed to noise. This rigidity limits the usefulness of Melissa as a tool to investigate intrinsic stochasticity in methylation at the single locus level. Relaxing the modelling assumptions to accommodate methylation variability in Melissa is an interesting topic for future research.

Another area where Melissa could be fruitfully applied is the integrative study of multiple high-throughput features in single cells. Kapourani and Sanguinetti (2016) showed that features extracted from methylation profiles could be effectively used to predict gene expression in bulk experiments. With the advent of novel technologies measuring gene expression and multiple epigenomic features in individual cells (Clark *et al*., 2018), interpretable Bayesian models like Melissa are likely to play an important role in furthering our understanding of epigenetic control of gene expression in single cells.

## 4 Methods

### 4.1 Modelling DNA methylation profiles

In order to provide spatial smoothing of the methylation profiles at specific loci, we adapt a generalised linear model of basis function regression proposed recently in Kapourani and Sanguinetti (2016). The basic idea is as follows: the methylation profile associated to a genomic region *M* is defined as a (latent) function *f*: *M* → (0,1) which takes as input the genomic coordinate along the region and returns the propensity for that locus to be methylated. In order to enforce spatial smoothness, and to obtain a compact representation for this function in terms of interpretable features, we represent the profile function as a linear combination of basis functions

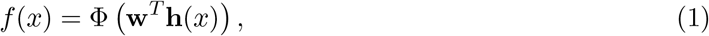

where **x** ≡ **h**(*x*) are the basis functions (here we consider radial basis functions (RBFs)), 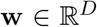 represent the regression coefficients, and Φ is the inverse probit function (Gaussian cumulative distribution function) needed in order to map the function output to the (0,1) interval. The latent function is observed at specific loci through a noise model which encapsulates the experimental technology; in Kapourani and Sanguinetti (2016), a Binomial observation model was proposed to handle bulk bisulfite sequencing data, and an efficient maximum likelihood strategy was proposed.

For single-cell methylation data, methylation of individual CpG sites can be naturally modelled using a Bernoulli observation model, since for the majority of covered sites we have binary CpG methylation states. To account for the inherent noise measurements and the limited CpG coverage, we also reformulate the model in a Bayesian framework. The Bayesian probit regression model for a single CpG site then becomes

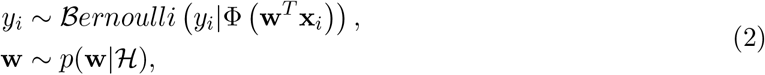

where *y_i_* denotes the CpG methylation state and 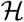 are (hyper)-parameters of the prior. Performing inference for this model in the Bayesian framework is complicated by the fact that no conjugate prior 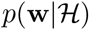 exists for the parameters of the probit regression model. The model can be made amenable to Bayesian estimation thanks to a data augmentation strategy originally proposed by Albert and Chib (1993). This strategy consists of introducing an additional auxiliary latent variable *z_i_*, which has a Gaussian distribution conditioned on the input **w**^*T*^**x**_*i*_. The augmented model has the hierarchical structure shown in Fig. 6, where *y_i_* is now deterministic conditional on the sign of the latent variable *z_i_*. Hence, our original problem becomes a missing data problem where we have a linear regression model on the latent variables *z_i_* and the observations *y_i_* are incomplete since we only observe whether *z_i_* > 0 or *z_i_* ≤ 0. Now we introduce a conjugate Gaussian prior over the parameters 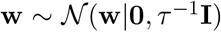, where the hyper-parameter *τ* controlling the precision of the Gaussian prior is assumed to follow a Gamma distribution. This reduces the necessary conditional distributions to a tractable form as either Gaussian, Gamma or one-dimensional truncated Gaussian distributions.

**Figure 6:**
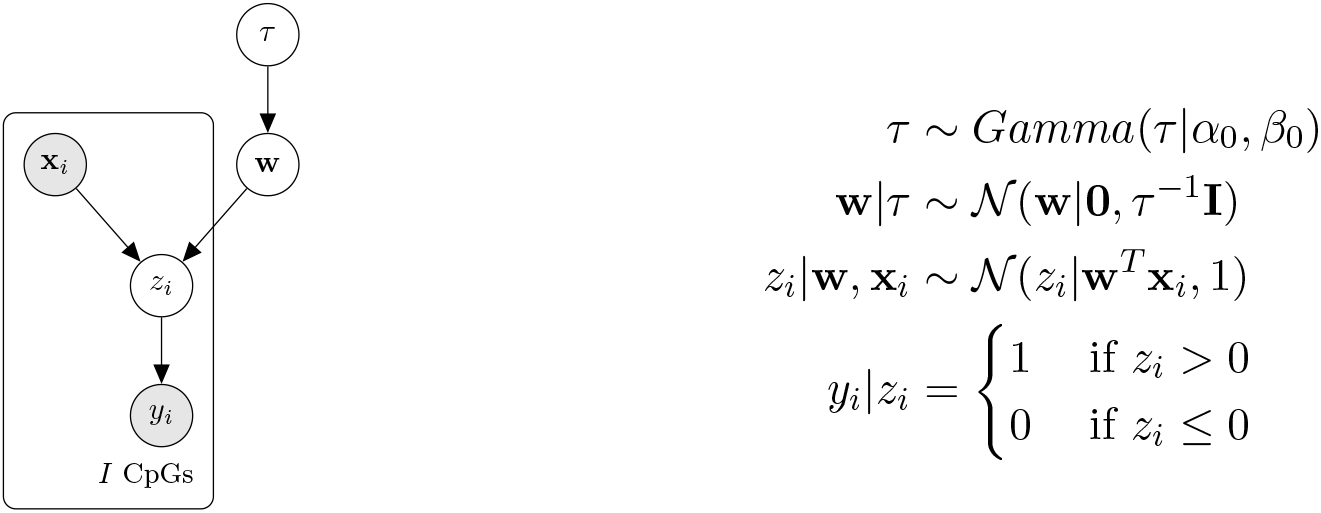
Probabilistic graphical representation of the Bayesian probit regression model.

### 4.2 Melissa model

Local smoothing certainly helps in dealing with data sparsity, however in our experience the coverage in scBS-seq data is insufficient to infer informative methylation profiles at many genomic regions. We therefore exploit the population structure of the experimental design to share and transfer information across cells.

Assume that we have *N*(*n* = 1,…, *N*) cells and for each cell we have a common set of *M*(*m* = 1,…,*M*) genomic regions, for example promoters, and we are interested in both partitioning the cells in *K* clusters and inferring the methylation profiles for each genomic region. To do so, we use a finite Dirichlet mixture model (FDMM) (McLachlan and Peel, 2004), where we assume that the methylation profile of the *m^th^* region for each cell *n* is drawn from a mixture distribution with *K* components (where *K* < *N*). This way cells belonging to the same cluster will share the same methylation profile, although profiles will still differ across genomic regions. Let *c_n_* be a latent variable comprising a 1-of-K binary vector with elements *c_nk_* representing the component that is responsible for cell *n*, and *π_k_* be the probability that a cell belongs to cluster *k*, i.e. *π_k_* = *p*(*c_n_* = *k*).

The conditional distribution of **C** given ***π*** is

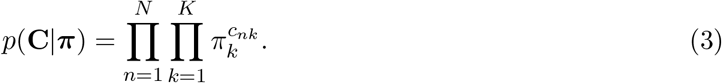

Considering the FDMM as a generative model, the latent variables **c**_*n*_ will generate the latent observations 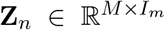, which in turn will generate the binary observations 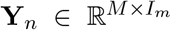 depending on the sign of **Z**_*n*_, as explained in the previous section. The conditional distribution of the data (**Z, Y**), given the latent variables **C** and the component parameters **W** becomes

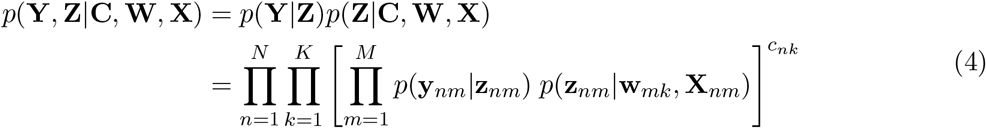

where

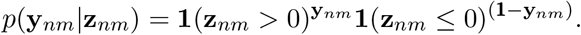

To complete the model we introduce priors over the parameters. We choose a Dirichlet distribution over the mixing proportions ***π***

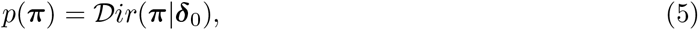

where for symmetry we choose the same parameter *δ*_0_*k*__ for each of the mixture components. We introduce an independent Gaussian prior over the coefficients W as we did in the Bayesian probit regression model

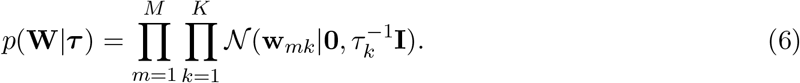

Finally, we introduce a prior distribution for the (hyper)-parameter ***τ***, and assume that each cluster has its own precision parameter

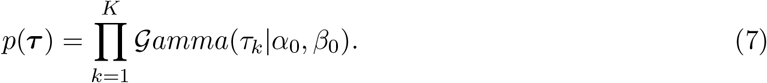

Having defined our model, we can now write the joint distribution over the observed and latent variables

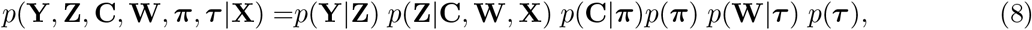

where the factorisation corresponds to the probabilistic graphical model shown in Fig. 7.

**Figure 7:**
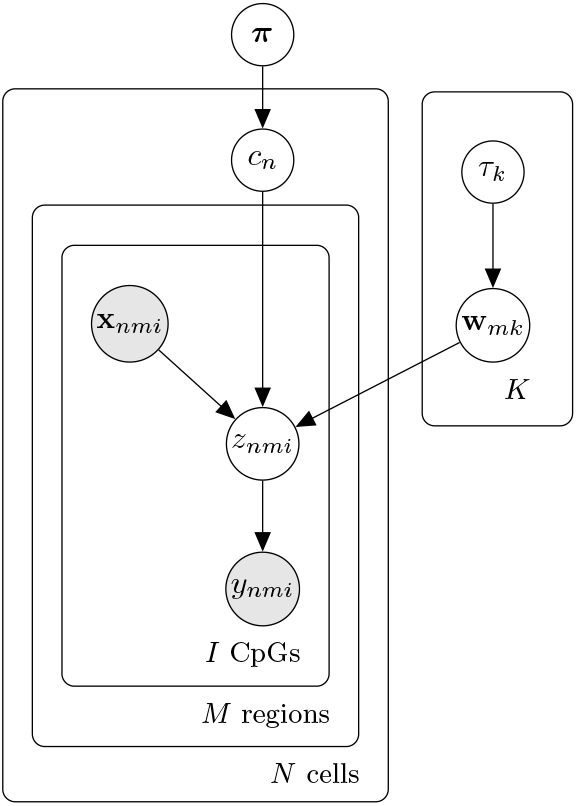
Probabilistic graphical representation of the Melissa model.

The standard approach now would be to compute the posterior distribution of the latent variables given the observed data *p*(**Z, C, W**, ***π, τ***|**Y, X**). However, exact computation of the posterior distribution is not analytically tractable and we have to resort to approximate techniques. We use mean-field variational inference (Blei *et al*., 2017) that exploits a factorisation assumption to achieve an efficient deterministic approximation to the posterior distribution.

#### Variational Inference

The most common method for performing approximate Bayesian inference for models with many parameters is Markov Chain Monte Carlo (MCMC) (Gelfand and Smith, 1990). However, frequently sampling methods require considerable computational resources and do not scale to perform genome-wide analysis on hundreds or thousands of single cells; variational methods can provide an efficient, approximate solution with better scalability in this case (see “Results” section for a comparison between Gibbs sampling and variational inference for this model). Besides the computational advantages, the deterministic nature of the variational inference machinery makes it easier to assess convergence compared to MCMC methods (Beal, 2003).

Briefly, in mean-field variational inference the intractable posterior distribution of the latent variables *p*(***θ***|**X**) is approximated by a factorized distribution *q*(***θ***) = Π_*i*_ *q_i_*(***θ***_*i*_), where ***θ*** denotes the latent variables and **X** the observed variables. Then we search over the space of approximating distributions to find the distribution with the minimum Kullback-Leibler 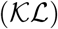 divergence with the actual posterior

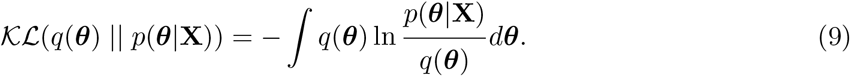

The 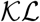 divergence can then be minimised by performing a free form minimisation over the *q_i_*(***θ***_*i*_) leading to the following update equation

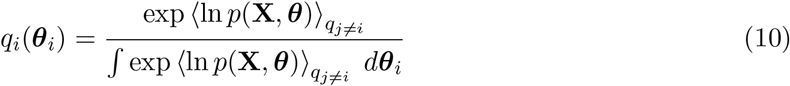

where 〈·〉*_q_j≠i__* denotes an expectation with respect to the distributions *q_j_*(***θ***_*j*_) for all *j* ≠ *i*. We can apply this approach to our probabilistic model and take the approximating distribution to factorise over the latent variables

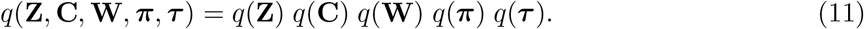

Applying Eq. (10) to our model, we obtain the following solutions for the optimised factors of the variational posterior

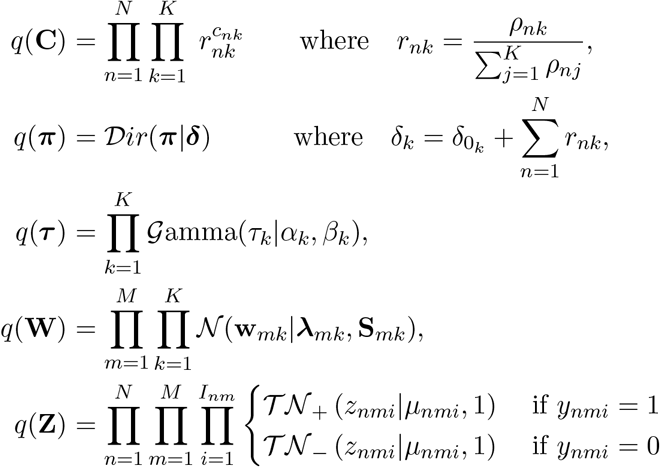

where

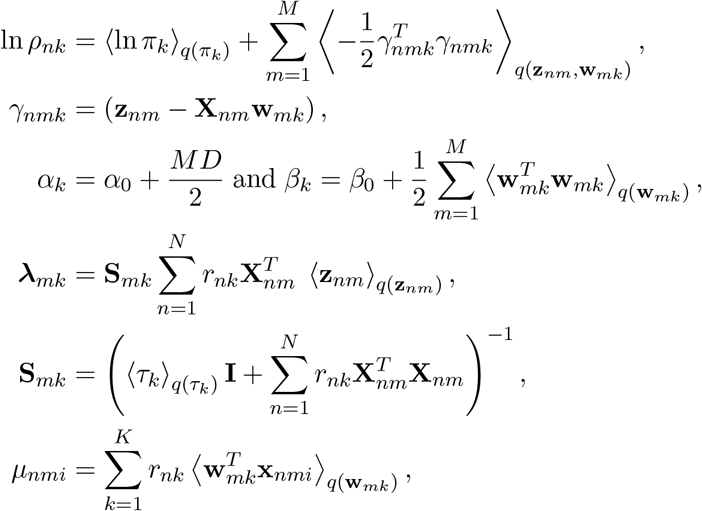

and 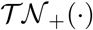 denotes the normal distribution truncated on the left tail to zero to contain only positive values, and 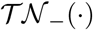 denotes the normal distribution truncated on the right tail to zero to contain only negative values. Detailed derivations are available in Additional file 1: Section 1.

#### Predictive density and model selection

Given an approximate posterior distribution, we are in the position to predict the methylation level at unobserved CpG sites. The predictive density of a new observation **y**_*_, which is associated with latent variables **c**_*_, **z**_*_ and covariates **X**_*_, is given by

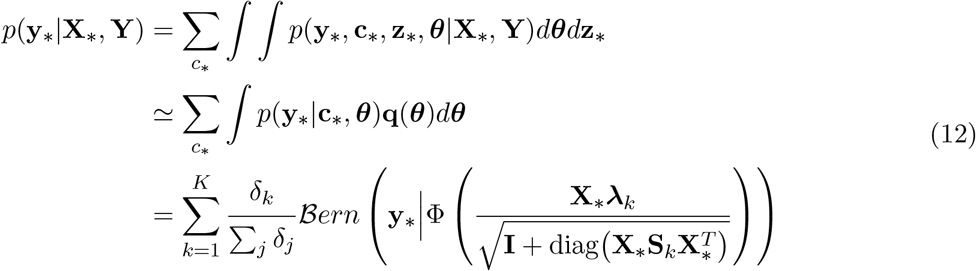

where we collectively denote as ***θ*** the relevant parameters being marginalised.

One of the most appealing aspects of variational approximations within mixture models is the possibility of directly performing model selection, i.e. determining the number of clusters, within the optimisation procedure. It has been repeatedly observed (Corduneanu and Bishop, 2001) that, when fitting variationally a mixture model with a large number of components, the variational procedure will prune away components with no support in the data, hence effectively determining an appropriate number of clusters in an automatic fashion. We can gain some intuition as to why this happens in the following way. We can rewrite the 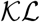 divergence as

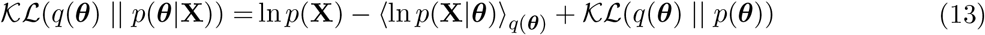

where ln *p*(**X**) can be ignored since is constant with respect to *q*(***θ***). To minimize this objective function the variational approximation will both try to increase the expected log likelihood of the data ln *p*(**X**|***θ***) while minimizing its 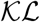 divergence with the prior distribution p(***θ***). Hence, using variational Bayes we have an automatic trade-off between fitting the data and model complexity (Bishop, 2006); giving the possibility to automatically determine the number of clusters without resorting to cross-validation techniques.

### 4.3 Assessing Melissa via a simulation study

To generate realistic simulated single-cell methylation data, we first used the BPRMeth package (Kapourani and Sanguinetti, 2018) to infer five prototypical methylation profiles from the GM12878 lymphoblastoid cell line. The bulk BS-seq data for the GM12878 cell line are publicly available from the ENCODE project (Dunham *et al*., 2012). Based on these profiles we simulated single cell methylation data (i.e. binary CpG methylation states) for M = 100 genomic regions, where each CpG was generated by sampling from a Bernoulli distribution with probability of success given by the latent function evaluation at the specific site. This process can be thought of as generating methylation data for a specific single cell. Next, we generated K = 4 cell sub-populations by randomly shuffling the genomic regions across clusters, so now each cell sub-population has its own methylome landscape. In total we generated N = 200 cells, with the following cell sub-population proportions: 40%, 25%, 20% and 15%. Finally, to account for different levels of similarity between cell sub-populations, we simulated 11 different datasets by varying the proportion of similar genomic regions between clusters. The R scripts for this simulation study are publicly available on the Melissa repository.

### 4.4 scBS-seq data and preprocessing

Two mouse embryonic stem cells (ESCs) datasets were used to validate the performance of the Melissa model. The first dataset presented in Angermueller *et al*. (2016), after quality assessment, consisted of 75 single cells out of which 14 cells were cultured in 2i medium (*2i ESCs*) and the remaining 61 cells were cultured in serum conditions (*serum ESCs*). The Bismark (Krueger and Andrews, 2011) processed data, with reads mapped to the GRCm38 mouse genome, were downloaded from the Gene Expression Omnibus under accession GSE74535. The second dataset (Smallwood *et al*., 2014) contained 32 cells out of which 12 cells were *2i ESCs* and the remaining 20 cells were *serum, ESCs* and the Bismark processed data, with reads mapped to the GRCm38 mouse genome, are publicly available under accession number GSE56879. For both datasets, binary single-base-pair CpG methylation states were obtained from the ratio of methylated read counts to total read counts.

The BPRMeth package (Kapourani and Sanguinetti, 2018) was then used to read single cell methylation data and for each cell create genomic methylation regions that would be used as input to Melissa. We considered six different genomic contexts where we applied Melissa: protein coding promoters with varying genomic windows: ±1.5kb, ±2.5kb and ±5kb around transcription start sites (TSS), active enhancers, super enhancers and Nanog regulatory regions. Due to the sparse CpG coverage, for the three genomic contexts except promoters we filtered loci with smaller than 1kb annotation length and specifically for Nanog regions we took a window of ±2.5kb around the centre of the genomic annotation. In addition, we only considered regions that were covered in at least 50% of the cells with a minimum coverage of 10 CpGs and had between cell variability; the rationale being that homogeneous regions across cells do not provide additional information for identifying cell sub-populations. We run the model with *K* = 6 and *K* = 5 clusters for the Angermueller et al. (2016) and Smallwood et al. (2014) datasets, respectively, and we use a broad prior over the model parameters.

### 4.5 Performance evaluation

To assess model performance across all genomic contexts, we partition the data and use 50% of the CpGs in each cell and region for training set and the remaining 50% as test set (except DeepCpG, see below). The prediction performance of all competing models, except DeepCpG, was evaluated on imputing all missing CpG states in a given region at once. For computing binary evaluation metrics, such as F-measure, predicted probabilities above 0.5 were set to one and rounded to zero otherwise.

**F-measure** The F-measure or F_1_-score is the harmonic mean of precision and recall:

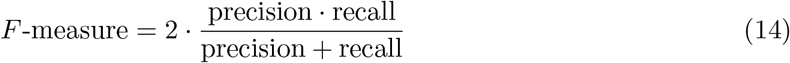

#### DeepCpG

The DeepCpG method takes a different imputation approach: it is trained on a specific set of chromosomes and predicts methylation states on the remaining chromosomes where it imputes each CpG site sequentially by using as input a set of neighbouring CpG sites. This approach makes it difficult to equally compare with the rival methods, since for each CpG the input features to DeepCpG are all the neighbouring sites, whereas the competing models have access to a subset of the data and they make predictions in one pass for the whole region. Since we only had access to CpG methylation data and to make it comparable with the considered methods, we trained the CpG module of DeepCpG (termed *DeepCpG CpG* in Angermueller *et al*. (2017)).

For the Angermueller *et al*. (2016) dataset, chromosomes 3 and 17 were used as training set, chromosomes 12 and 14 as validation set and the remaining chromosomes as test set. For the Smallwood *et al*. (2014) dataset, chromosomes 3, 17 and 19 were used as training set, chromosomes 12 and 14 as validation set and the remaining chromosomes as test set. The chosen chromosomes had at least 3 million CpGs used as training set; a sensible size for the DeepCpG model as suggested by the authors. A neighbourhood of K = 20 CpG sites to the left and the right for each target CpG was used as input to the model. During testing time, even if a given genomic region did not contain at least 40 CpGs, the DeepCpG model used additional CpGs outside this window to predict methylation states; hence using more information compared to the rival models. In total the DeepCpG model took around four days per dataset for training and prediction on a cluster equipped with NVIDIA Tesla K40ms GPUs.

**Adjusted Rand Index** The Adjusted Rand Index (ARI) is a measure of the similarity between two data clusterings:

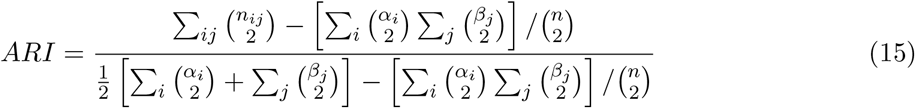

## Declarations

### Acknowledgements

We thank Oliver Stegle, Ricard Argelaguet, Michalis Michaelides, Duncan Sproul and Stephen Clark for valuable comments and discussion.

### Software availability

Melissa is implemented in the R programming language and is publicly available from the following repository: https://github.com/andreaskapou/Melissa, which also contains all the scripts for reproducing the manuscript results. The method will shortly be available as a Bioconductor package.

